# FiberNET: An ensemble deep learning framework for clustering white matter fibers

**DOI:** 10.1101/141036

**Authors:** Vikash Gupta, Sophia I. Thomopoulos, Faisal M. Rashid, Paul M. Thompson

## Abstract

White matter tracts are commonly analyzed in studies of micro-structural integrity and anatomical connectivity in the brain. Over the last decade, it has been an open problem as to how best to cluster white matter fibers, extracted from whole-brain tractography, into anatomically meaningful groups. Some existing techniques use region of interest (ROI) based clustering, atlas-based labeling, or unsupervised spectral clustering. ROI-based clustering is popular for analyzing anatomical connectivity among a set of ROIs, but it does not always partition the brain into recognizable fiber bundles. Here we propose an approach using convolutional neural networks (CNNs) to learn shape features of the fiber bundles, which are then exploited to cluster white matter fibers. To achieve such clustering, we first need to re-parameterize the fibers in an intrinsic space. The clustering is performed in induced parameterized coordinates. To our knowledge, this is one of the first approaches for fiber clustering using deep learning techniques. The results show strong accuracy - on a par with or better than other state-of-the-art methods.

## 1 Introduction

White matter fibers are important structures in the brain, connecting its various components, and vulnerable to breakdown in a variety of brain diseases. Studying white matter (WM) fiber bundles brings new insight into disease progression, and into the structural network supporting communication in the brain. The brain’s neural pathways - or fiber tracts - have complex individual variations in geometry, and they are interspersed with each other - which makes it very difficult to cluster them into anatomically meaningful groups or units for further statistical analysis. One commonly used clustering method [12] uses manual ROI delineation on the images of fractional anisotropy (FA), a scalar metric derived from diffusion-weighted MRI. These regions can be used to seed whole-brain fiber tractography, and the set of resulting curves - or streamlines - is then grouped into white matter bundles using spectral clustering. One method uses Hausdorff’s distance [4] as a distance metric between two fibers, prior to hierarchical clustering based on the distances between fibers. Recently, several unsupervised clustering methods [11, 15, 20] have been proposed. Similar flavors of unsupervised techniques and an understanding of hierarchical clustering of fibers in video analytics have been explored in [10]. We have also explored the unsupervised learning procedures like support vector machines (SVM) and spectral clustering in [6,7]. Though mathematically elegant, these methods come with a baggage of assumptions and thus with their own limitations. However, with the increasing amount of data, we can make a completely data driven clustering algorithm free of any assumptions using convolution neural network (CNN) framework. Though, texture based neural networks have been used even in the early years of brain imaging [14], they were not used for white matter clustering. Recently, CNNs have been extensively used in the computer vision community for object detection, clustering and segmentation [17]. Deep learning methods offer some attractive properties. The most important is *automatic feature selection*. A well designed CNN should be able to extract the most discriminative features to achieve a given task. Another attractive feature of this framework is *re-usability* of the model. A model trained on one dataset can be used for classifying another dataset; we can test how well this “transfer learning” approach works, and what factors in the input data or method make it perform best. Another benefit of such a system is the *scalability*. We can start with a small number of training datasets, and the network will automatically improve over time as we add increasing amounts of data to train the model. To use the CNN framework, we introduce a volumetric parameterization technique to transform the brain into a topologically equivalent spherical domain. In computational anatomy, many algorithms have been devoted for surface parameterization [5, 18]. Surface parameterization may be sufficient for analyzing surface geometry. However, it falls short when there is significant information contained inside the shape under consideration (e.g., WM fibers in the brain). Following the “sphere carving” work by Wang and colleagues [19] in this paper, we developed a novel method that parameterizes the entire volume of the brain and every structure contained in it. We then use the parameterized coordinates of the tracts to cluster the white matter fibers.

## 2 Harmonic Function

We parameterize the 3D volume using a potential function *ϕ* with harmonic property. A function with the harmonic property is a *C*^2^ continuous function that satisfies Laplace’s equation. Harmonic functions can be used to establish a bijective mapping between the brain and the topologically equivalent spherical shape. If *ϕ* : *U*→*R*, where *U*⊆*R^n^* is some domain and *ϕ* is some function defined over *U*, the function *ϕ* is harmonic if its Laplacian vanishes over *U*, i.e., ∇^2^*ϕ* = 0. In terms of Cartesian coordinate system, we can write

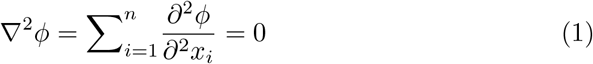

where *x_i_* is the *i^th^* Cartesian coordinate and *n* is the number of dimensions of the shape under study (here, 3).

### 2.1 Defining a shape-center

Ideally, the shape-center should be located at approximately the same anatomical location. This particular point is located at (98, 113, 111) in the standard MNI template. The corresponding point among the subjects is located using a linear registration process. The B_0_, T1 images and the MNI templates are rigidly registered in the same order. The transformations are combined and inverted. When applied to the aforementioned point, the inverse transformation outputs the shape-center in the subject space.

### 2.2 Boundary Conditions

We apply the Dirchlet and Neumann boundary conditions on the shape-center and the boundary surface (∂*U*), i.e., we assign the value of the function *ϕ* on all the boundary points and the shape-center to 1 and 0 respectively. These values remain unchanged across computation. All the remaining points inside the brain are assigned random values between 0 and 1 as the initial condition.

### 2.3 Potential Computation

An iterative finite difference scheme is used to solve the Laplace equations. If *ϕ*(*x, y, z*) is a harmonic function, its second derivative is computed using the Taylor’s series expansion and using the Laplace equation from 1 we have

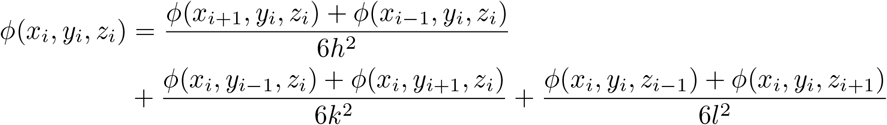

where *h, k* and *l* are the grid resolution in *x, y* and *z* directions respectively. The above potential values are computed until the maximum difference between two successive iterations is below a certain threshold *ζ* (< 10^−6^).

### 2.4 Computing potential-flow lines

Streamlines or the potential flow lines are orthogonal to the equi-potential surfaces created in the previous step. Each of the streamlines starts from the boundary points on the brain surface and progresses towards the designated shape-center. Each of these streamlines approaches the shape-center at unique angle(s), which remain constant along the streamline. This property is endowed by construction. The streamlines are computed by solving the following differential equation,

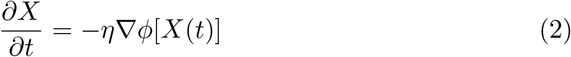

where *X* = [*x, y, z*]^*T*^ is the coordinate vector and *η* is the normalization constant. MATLAB’s *ode23* routine is used to solve the system of differential equations.

### 2.5 Parameterizing the Brain

Each streamline originating from each of the boundary points approaches the shape-center at a unique angle of approach. These angles remain constant along the streamlines. In case of three dimensional objects, the angle of approach is characterized by the elevation (*θ*) and the azimuthal (*ψ*) angles. The vector between the shape-center and the end point of the streamline is calculated. The angles are calculated using the Cartesian to spherical coordinate transformation

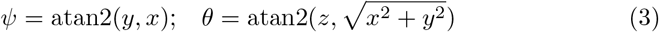

The streamlines intersect the equipotential surfaces at right angles (see figure 1). Each point of intersection generates a tuple [*ϕ, θ, ψ*]^*T*^ for the corresponding Cartesian coordinates [*x, y, z*]^*T*^.

**Fig. 1.**
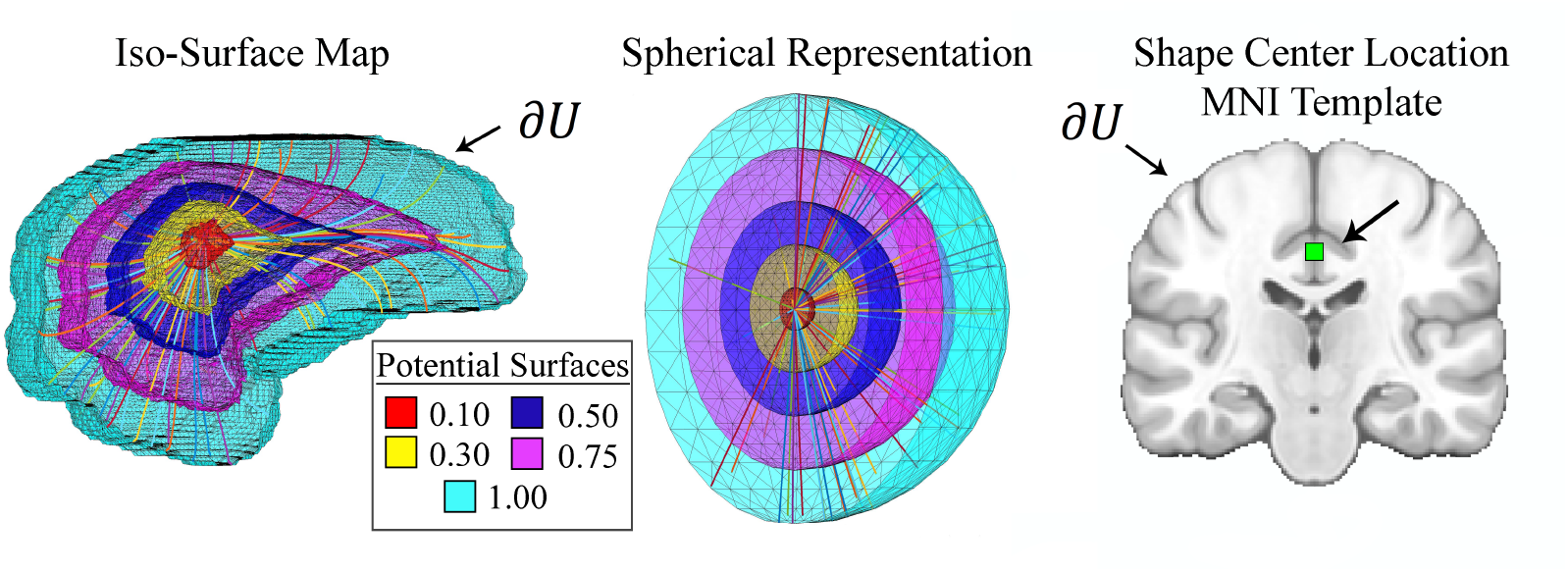
**Left:** 3D view of different equipotential surfaces are shown. The streamlines lines emanating from the surface approach the shape-center at unique polar and azimuthal angles. These *angles of approach* remain constant along the streamlines and intersect the surfaces at right angles. **Middle:** The equivalent spherical shape. The streamlines are radial lines emanating from the surface. The different colors show the potential value of equipotential surfaces. **Right:** Location of shape-center on the MNI template.

## 3 Mapping the White Matter Fibers

After the whole brain is parameterized as above, each fiber tract is mapped to the new coordinate system, i.e., in the spherical space. At this stage, we have a bijective mapping between the Cartesian coordinates of every voxel in the brain and the newly computed coordinate system. A KD-tree accessor is built using the native brain coordinates for *ϕ, θ* and *ψ*. For every point on the fiber streamline, the algorithm searches for ten neighborhood points and computes a weighted average to get the corresponding coordinate in the target domain. This process establishes the mapping of fibers in the target spherical domain.

## 4 Data pre-processing and fiber tracking

Each of the diffusion images has 46 volumes acquired with 5 T2-weighted B0 volumes and 41 diffusion-weighted volumes with voxel size of 1.36 × 1.36 × 2.7 mm^3^. The scans are acquired using a GE 3.0T scanner, using with echo planar imaging with parameters: TR/TE = 9050/62 ms. The raw DWI volumes are aligned to the first b_0_ image using FSL’s *eddy_correct* [16] to correct for head motion and eddy current distortions. The corresponding T1 image is skull stripped and a brain-mask was calculated. The B0 and T1 images are registered using an affine registration. The resulting inverse transformation is used to transfer the mask was transferred to the diffusion image space. Diffusion tensors and whole brain probabilistic tractography [13] are computed using the Camino toolkit [3]. Voxels with fractional anisotropy (FA) values greater than 0.2 were chosen as seed points. The maximum fiber turning angle was set to 60°/voxel, and tracing stopped at any voxel with FA < 0.2.

### 4.1 Data Augmentation

One challenge in multi-class classification problems like the one presented here is the uneven training samples for each class. Ideally, the training samples should contain an even number of samples to avoid bias against any particular class. However, it is natural to have an unbalanced group as some fiber bundles will be thicker and some fiber bundles will be very narrow. In order to have an even distribution of the training samples, we create copies of the original data set by convolving with a three 1-D Gaussian filters along *x, y* and *z* axes to create noisy variants of the tracts in the training dataset. In addition, the order of points in the tracts is flipped. This process makes the training process invariant to the order of points in the tracts.

## 5 Network architecture

The augmented data is mapped to the spherical domain as described in section 3. For our task the network architecture is presented below. We experimented with multiple kernel sizes and layers and present the most successful architecture. The network contains two convolutional layers with 32 and 64 feature maps. Feature maps define the number of filters used in each layer, and their sizes determine the receptive field. A large receptive field can acquire higher-order spatial information while the trade-off is the increase in the number of parameters. The non-linearity inducing functions used are rectified linear units (ReLU) and the hyperbolic tangent function (tanh). To prevent over-fitting we use 80% dropout [8], which randomly switches off neuronal units in each layer thereby reducing their influence at any particular iteration during back-propagation. Finally, the last layer is a ‘softmax’ layer with 17 outputs for 17 classes. In the present work, the deep learning is implemented using the TensorFlow version r0.11 [1].

**Fig. 2.**
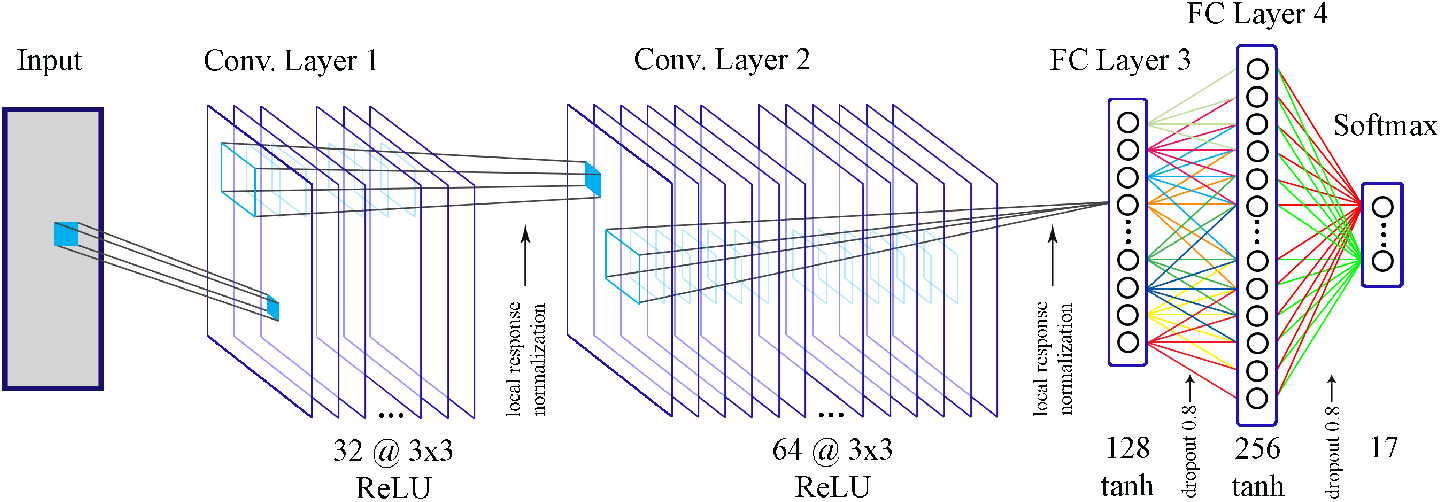
Each fiber in the input layer is a 50 × 3 matrix. There are two convolutional layers of size 32 and 64 respectively. Each layer contains convolutional kernels of size 3 × 3. The last convolutional layer is followed by two fully connected (FC) layers of size 128 and 256 followed by a softmax (output) layer.

### 5.1 Training data and majority voting

Out of 96 subjects, manual segmentation is performed on four randomly selected subjects. These four subjects act as the training data set. Each subject’s tractography data is manually clustered into 17 anatomically relevant fiber bundles. We use the manual tract segmentation on each of the four subjects used as the training data for the method. Then we perform a leave one out cross-validation (LOOCV) procedure. Another method to improve the accuracy of the method was to use bootstrap aggregation [2]. We sample our data with replacements for 600,000 fibers, 20 times. In essence we fit 20 different models to the given dataset. For prediction, we used the majority voting procedure from all the different models for prediction.

## 6 Results

The LOOCV process proves the feasibility of our network architecture. Further, we compare our method to an existing automatic clustering algorithm, autoMATE [9]. The reliability can be assessed using the confusion matrix. In Figure 3, we show two matrices that compare the accuracy of predictions. The diagonal and the off-diagonal values represent the true positive and the false positive rates of prediction. On the left, we show the mean prediction accuracy on the 4 cross-validation dataset. The prediction labels are compared against manual segmentation. On the right, we show a similar mean matrix for all the 92 subjects in the dataset, when compared to autoMATE [9]. One of the drawbacks of autoMATE is that it is very conservative, in the sense that it will not misclassify tracts, but it is quite likely that it will miss out on true positives. The presented method has no such bias. In Figure 4, the left panel shows a comparison of FiberNET with manual segmentation. On the right, a comparison between FiberNET and autoMATE is presented. As we can see, FiberNET mis-classifies certain tracts when compared against a random test subject from the dataset. However, we would like to argue that the CNN based method is much more flexible and gives us an opportunity to search for a better architecture that could possibly address the misclassification problem as it happened in the computer vision community, though challenges like ImageNET.

**Fig. 3.**
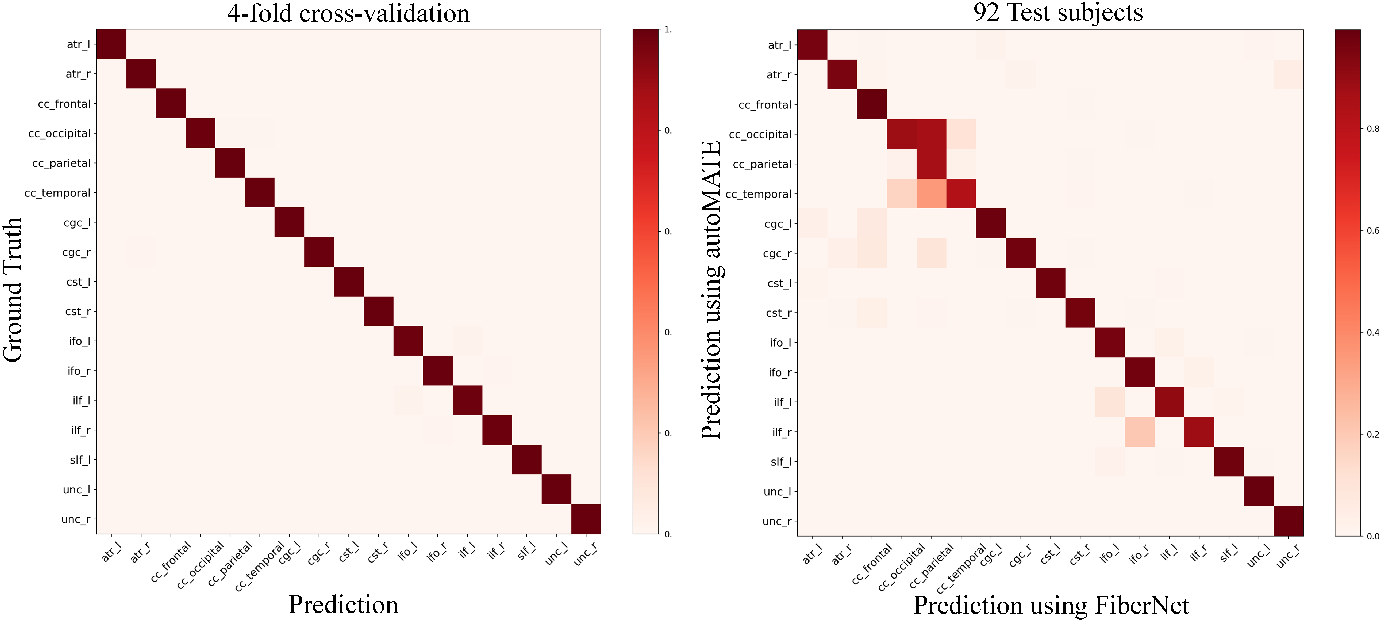
The confusion matrices show the classification accuracy of FiberNET. **Left:** Validation set of 4 subjects. **Right:** 92 subjects in the dataset when compared to autoMATE [9]

**Fig. 4.**
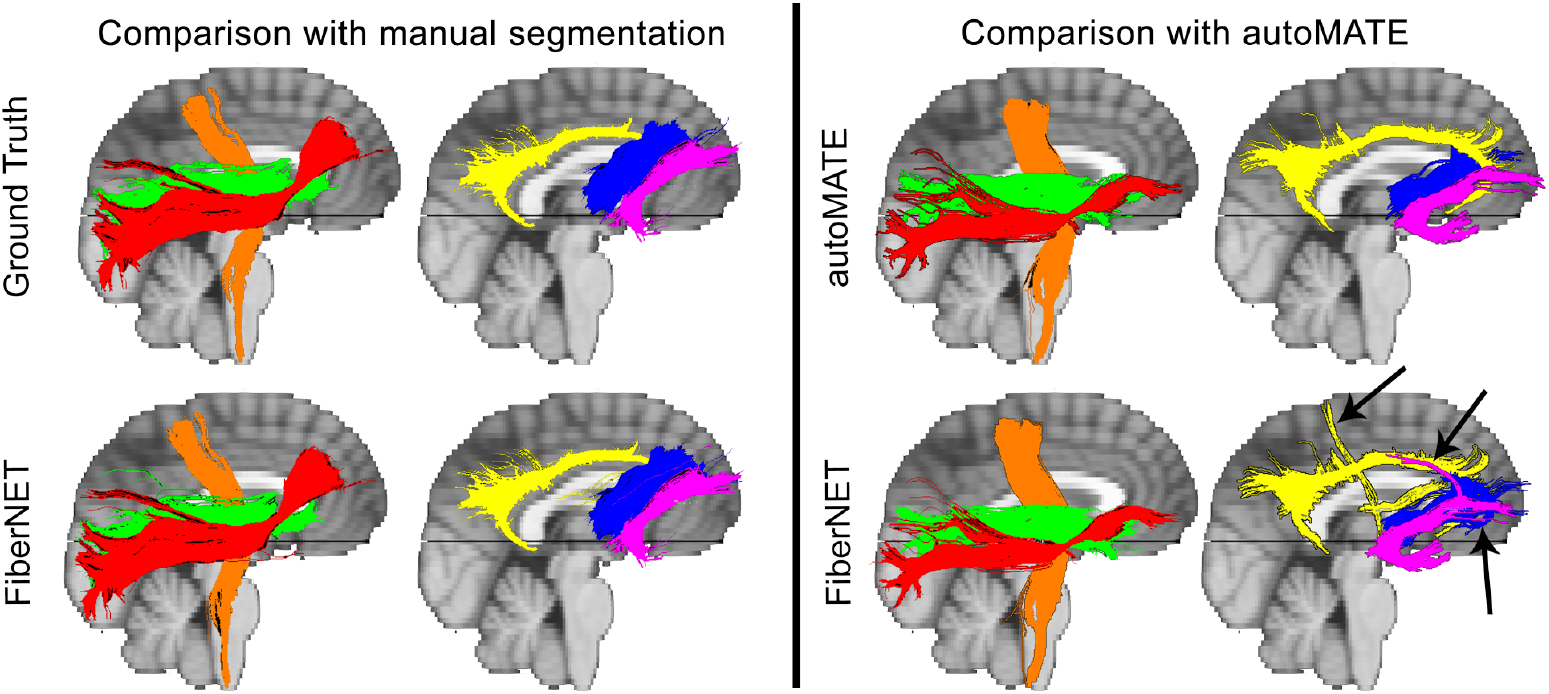
Comparison of ground truth (top row) versus predicted clusters (bottom row) on a validation subject (left) and a random test subject (right). We present here six representative tracts to show examples of good and average clustering; the right inferior fronto-occipital fasciculus (red), right cortico-spinal tract (orange), right inferior longitudinal fasciculus (green), right cingulum (yellow), right uncinate fasciculus (pink), and right anterior thalamic radiation (blue). The arrows show the mis-classified tracts.

### 6.1 Conclusions

In this paper, we presented an ensembled deep learning approach to cluster white matter fibers into anatomically meaningful fiber bundles. We have shown for the first time a reliable method to apply deep learning approaches to the fiber clustering problem. Though our method is not 100% accurate in case of a few fiber bundles, this is understandable as they are equally difficult to cluster even manually by a neuroanatomy expert, because of the underlying complexity of the white matter structures. Nonetheless, this is a first step towards solving this clustering problem using a neural network. We also expect that the reliability of the method should increase with time as more datasets are added to the model, to better capture the spectrum of variability. This hypothesis is based on the near-perfect results we see in the computer vision community for the case of image classification problems. We also proposed a volumetric parameterization procedure to define an intrinsic coordinate system for the brain. We showed the efficacy of such parameterization in solving the white matter classification problem. We also believe that this parameterization method can be further explored for interesting brain imaging applications such as registration and ROI segmentation.

## 7 Appendix

The following table shows the abbreviations of all the 17 white matter fibers considered for clustering in this paper.

| Abbreviation | Fiber bundle name |
| --- | --- |
| atr_l | anterior thalamic radiation (left) |
| atr_r | anterior thalamic radiation (right) |
| cc_frontal | the corpus callosum section connecting frontal lobes |
| cc_occipital | corpus callosum section connecting occipital lobes |
| cc_parietal | corpus callosum section connecting parietal lobes |
| cc_temporal | corpus callosum section connecting temporal lobes |
| cgc_l | cingulate gyrus part of cingulum (left) |
| cgc_r | cingulate gyrus part of cingulum (right) |
| cst_l | corticospinal tract (left) |
| cst_r | corticospinal tract (right) |
| ifo_l | inferior fronto-occipital fasciculus (left) |
| ifo_r | inferior fronto-occipital fasciculus (right) |
| ilf_l | inferior longitudinal fasciculus (left) |
| ilf_r | inferior longitudinal fasciculus (right) |
| slf_l | superior longitudinal fasciculus (left) |
| unc_l | uncinate fasciculus (left) |
| unc_r | uncinate fasciculus (right) |

To corroborate our findings, we are presenting additional results. In figures 5 and 6, we compare our results to that of manual segmentation on the 4 training subjects. These results were obtained through a leave one out cross validation (LOOCV) procedure, i.e., the training data was comprised of 3 of the four subjects and the remaining subject was the validation subject. The cycle was repeated for all the 4 subjects. In Figure 7, we show a comparison between autoMATE [9] and the proposed method, FiberNET. The two subjects are chosen at random from the set of 92 subjects.

**Fig. 5.**
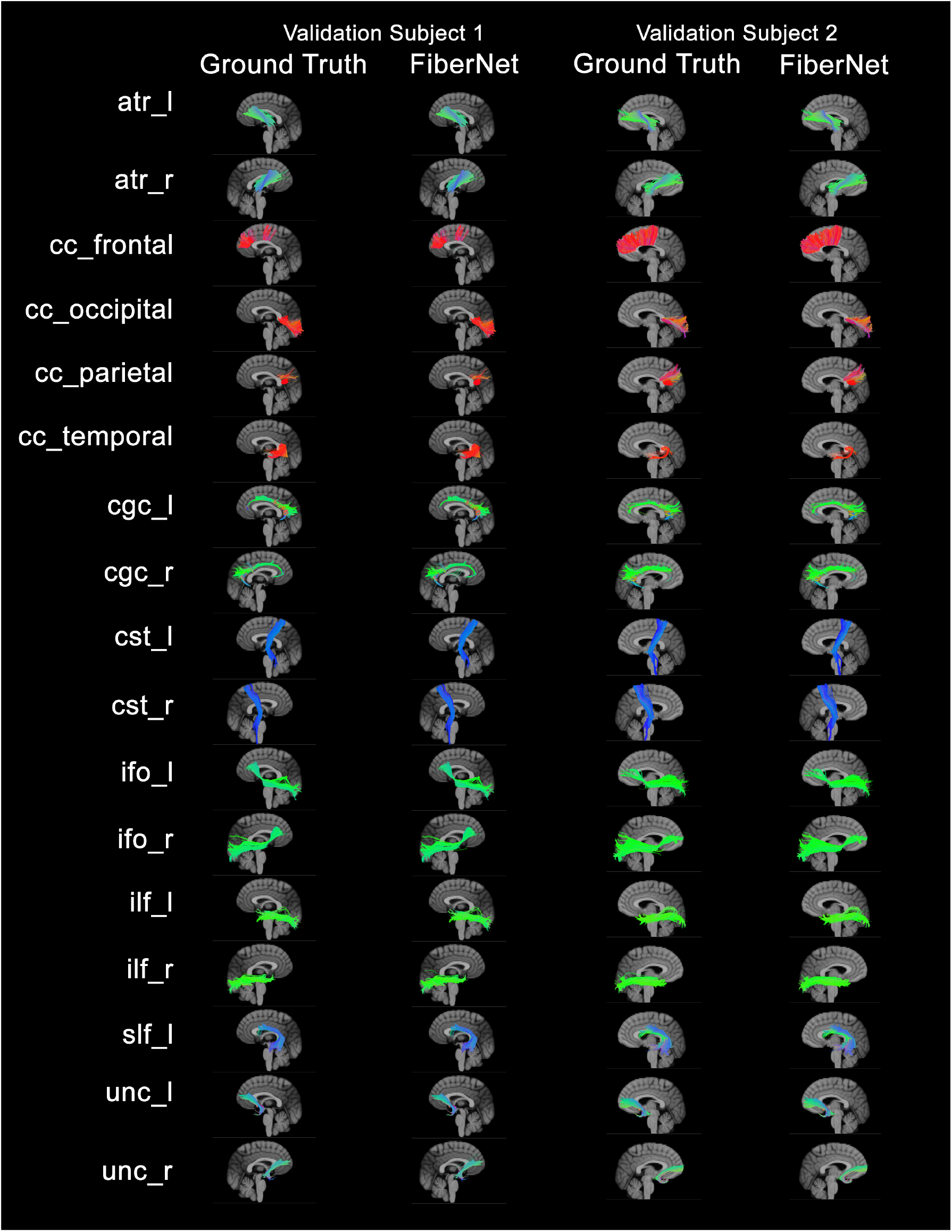
Comparison between ground truth and FiberNET prediction on 2 subjects from the four-fold cross validation (part 1 of 2)

**Fig. 6.**
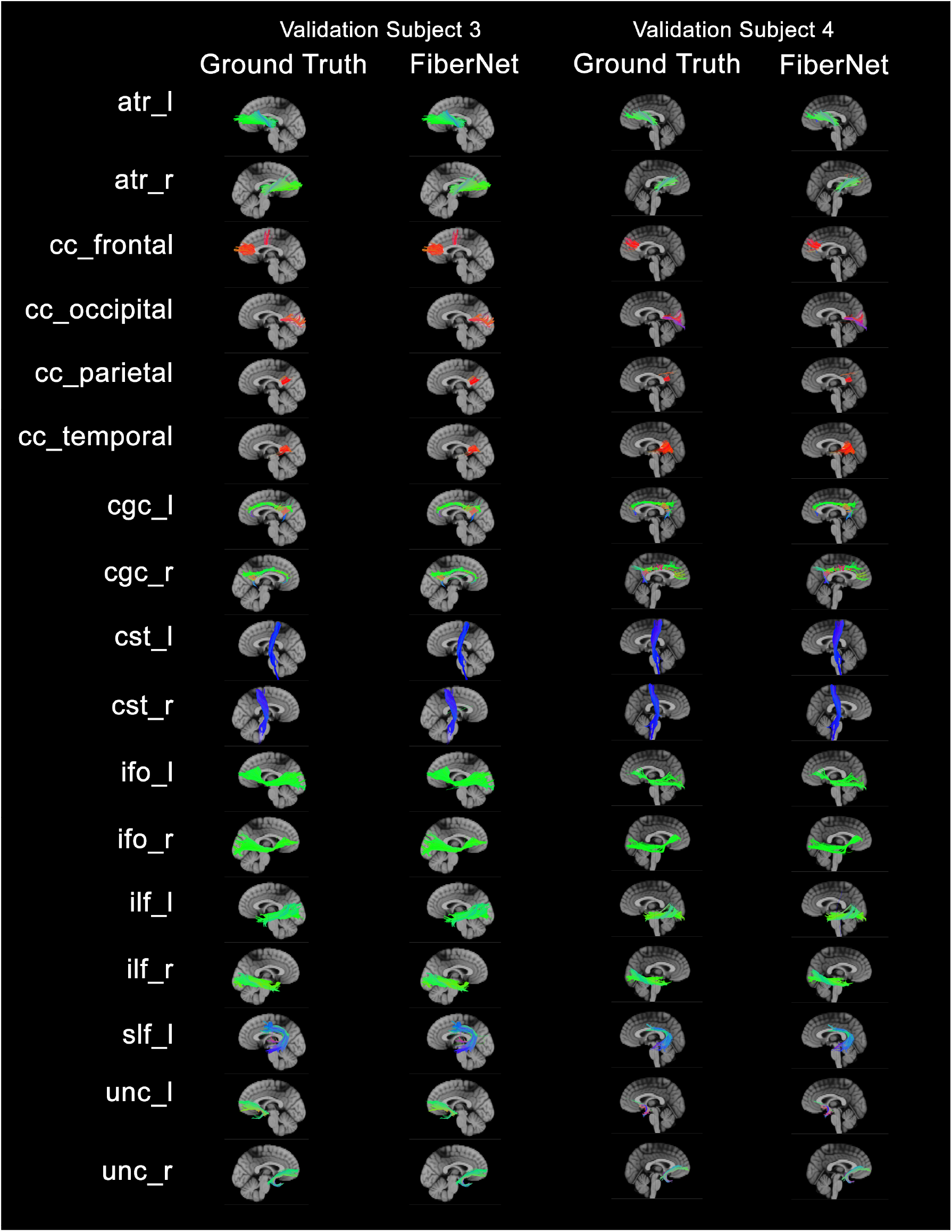
Comparison between ground truth and FiberNET prediction on 2 subjects from the four-fold cross validation (part 2 of 2)

**Fig. 7.**
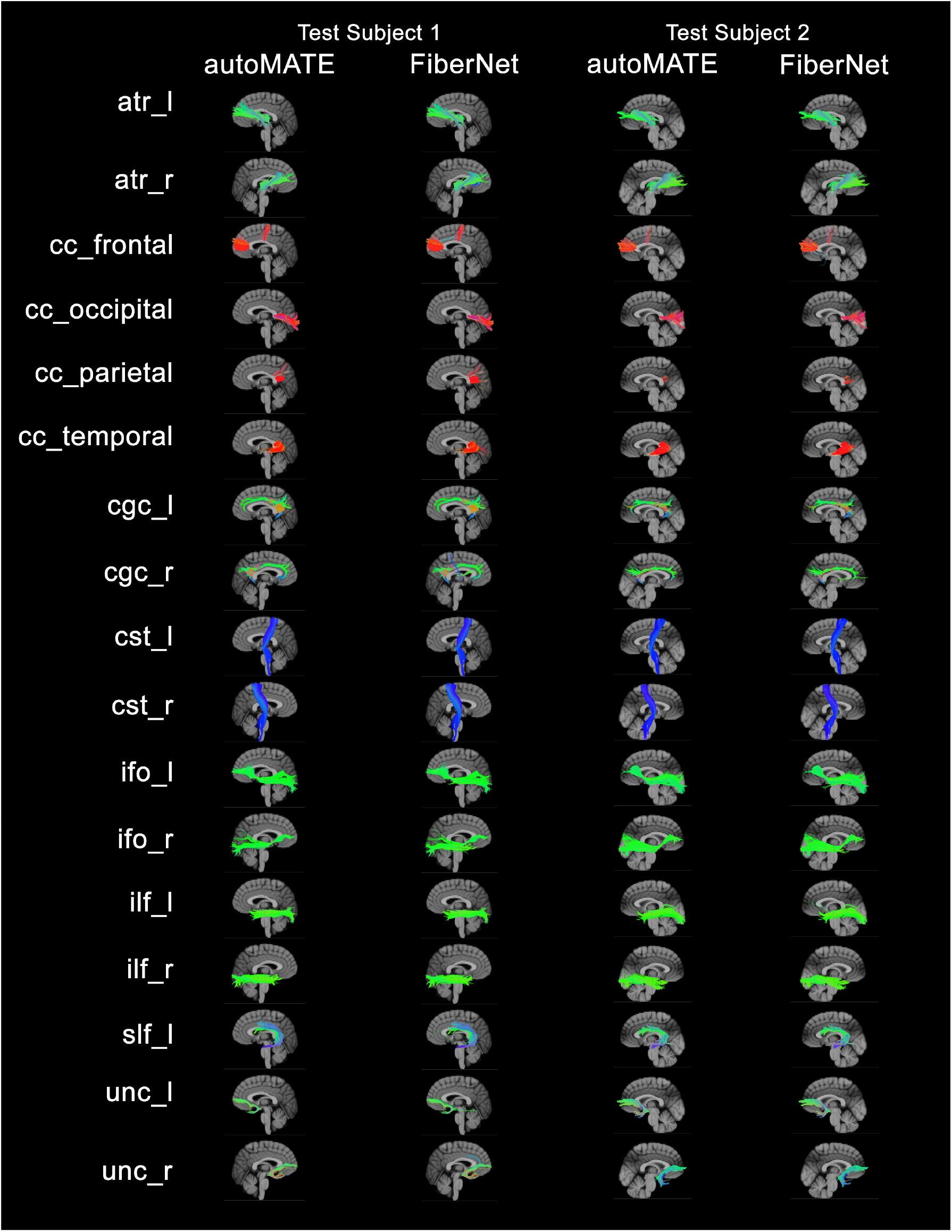
Comparison between autoMATE and FiberNET predictions on two random test subjects from a database of 92 subjects.

